# Cardiovirus-Mediated PKR Inhibition Results from Nucleocytoplasmic Trafficking Disruption

**DOI:** 10.1101/2025.08.01.668079

**Authors:** Romane Milcamps, Belén Lizcano-Perret, Fanny Wavreil, Marielle Lebrun, Chiara Aloise, Didier Vertommen, Gaëtan Herinckx, Frank J.M. van Kuppeveld, Catherine Sadzot, Thomas Michiels

**Affiliations:** Université catholique de Louvain, de Duve Institute, Brussels, Belgium; ULiege, GIGA-Immunobiology, laboratory of Virology and Immunology, Belgium; Utrecht University, Section of Virology, Division of Infectious Diseases and Immunology, Department of Biomolecular Health Sciences, Faculty of Veterinary Medicine, Utrecht, the Netherlands; MASSPROT platform, de Duve Institute, Université Catholique de Louvain, Brussels, Belgium

**Keywords:** PKR, EIF2AK2, nucleocytoplasmic traffic, nucleocytoplasmic trafficking disorder (NCTD), picornavirus, cardiovirus, nucleoli, double-stranded RNA, stress granules

## Abstract

Eukaryotic translation initiation factor 2 alpha kinase 2 (EIF2AK2), know as PKR, is a key antiviral kinase activated by double-stranded RNA (dsRNA) typically produced during viral replication. Upon activation, PKR phosphorylates eIF2α, leading to the inhibition of translation and viral replication. However, many viruses have evolved mechanisms to counteract PKR activity. In Cardioviruses, the Leader protein (L), a short peptide cleaved from the N-terminus of the viral polyprotein, not only inhibits PKR but also blocks interferon production and disrupts nucleocytoplasmic trafficking (NCT). L disrupts NCT by recruiting host RSK kinases to the nuclear pore complex (NPC), where RSK phosphorylates FG-nucleoporins, thereby impairing NCT. L mutations that affect NCT disruption also impact its ability to inhibit PKR, suggesting a mechanistic link. Recombinant TMEV and EMCV viruses designed to disrupt NCT through different mechanisms exhibited some extent of PKR inhibition, supporting the link between NCT disruption and PKR inhibition. Immunostaining and live-cell imaging revealed that L-induced NCT disruption redistributes a fraction of PKR to the nucleoli, where PKR remains inactive. This suggests that nucleolar sequestration contributes to PKR inhibition. Additionally, L-mediated NCT disruption leads to the release of nuclear RNA-binding proteins (nRBPs) into the cytosol, which may bind or modify viral dsRNA, further preventing PKR activation. Collectively, these results highlight nucleocytoplasmic trafficking as a critical regulatory mechanism governing PKR activation. Thus, beyond the specific action of cardiovirus L protein, our study reveals that interference with host nucleocytoplasmic transport can significantly impact the subcellular localization and functional regulation of immune effectors such as PKR.

**Author Summary:** Protein kinase R (PKR) is a crucial component of the host innate immune response. It is activated by double-stranded RNA (dsRNA) typically produced during viral replication and triggers a shutdown of mRNA translation. This antiviral mechanism limits viral propagation by inhibiting both host and viral protein synthesis. However, many viruses have developed mechanisms to inhibit PKR, allowing them to escape immune detection. PKR downregulation facilitates viral replication whereas uncontrolled PKR activation can lead to autoimmune disorders. Therefore, PKR activity must be tightly regulated to maintain immune homeostasis. Using recombinant viruses which target the nuclear pore complex, we show that nucleocytoplasmic trafficking of cellular components is critical for regulation of PKR activity. Infection of cells with Theiler’s murine encephalomyelitis virus triggers an efflux of nuclear RNA binding proteins which likely compete with PKR for dsRNA binding and thereby block PKR activity. Moreover, upon TMEV infection as well as during mitosis, PKR is detected in the nucleoli where it is thought to interact with structured RNAs without being activated. Our data highlight an important link between nucleocytoplasmic trafficking and PKR activity.

## Introduction

The integrated stress response (ISR) is a crucial signaling pathway conserved across mammalian species, activated by diverse stress stimuli. The central event of this process is the phosphorylation of the alpha subunit of the eukaryotic translation initiation factor 2 (eIF2α) by one of four members of the eIF2α kinase family (EIF2AK1-4) (1, 2). eIF2α phosphorylation leads to a global shutdown of protein synthesis while selectively inducing ATF4 translation and ATF4-dependent transcription of specific genes that collectively support the restoration of cellular homeostasis (3). EIF2AK1 (HRI) primarily senses oxidative stress, EIF2AK3 (PERK) responds to endoplasmic reticulum stress, and EIF2AK4 (GCN2) is activated by amino acid deprivation (1, 2). EIF2AK2, commonly known as PKR, is an interferon-induced protein kinase primarily activated by viral infection. PKR is a 551 amino acid-long cytoplasmic serine-threonine kinase harboring two amino-terminal double-stranded RNA-binding motifs (DRBMs) separated from a carboxy-terminal catalytic kinase domain (KD) by a flexible linker. In its inactive state, PKR adopts a closed conformation, where the second DRBM interacts tightly with the kinase domain, concealing the substrate-binding pocket and preventing its inadvertent activation (4). During viral infection, PKR is activated by binding double-stranded RNA (dsRNA) molecules, a highly immunogenic intermediate formed during viral replication. Upon dsRNA binding, PKR self-associates to form homodimers and undergoes autophosphorylations to achieve full activation (5). Phosphorylation of Thr446 and Thr451 within the activation loop is essential for PKR activation and can serve as a marker for monitoring its activation state (6). Once activated, PKR phosphorylates its substrate, eIF2α, at Ser51, increasing its affinity for eIF2B, a guanine nucleotide exchange factor for eIF2. This interaction inhibits the recycling of GDP-bound eIF2α, thereby preventing the formation of the translation preinitiation complex. Translation blockade inhibits both cellular and viral mRNA translation, leading to the rapid formation of stress granules (SG) (7). By preventing viral mRNA translation and by stimulating infected cell apoptosis (5), PKR acts as a major antiviral arm of the innate immune response. Given its critical role, numerous viruses have evolved mechanisms to block the PKR-mediated antiviral response (8).

Cardioviruses, which belong to the *Picornaviridae* family, are non-enveloped viruses with a genome consisting of single-stranded RNA molecule of positive polarity. The genus *Cardiovirus* includes encephalomyocarditis virus (EMCV), Theiler’s murine encephalomyelitis virus (TMEV), and human Saffold virus (SAFV). Upon delivery into the cell cytoplasm, the viral genome is directly translated into a single polyprotein through internal ribosome entry site (IRES)-mediated translation. The resulting polyprotein is then processed into structural and non-structural proteins, mainly by viral protease 3C. After the initial synthesis of viral proteins, viral RNA replication mediated by the viral 3D polymerase begins, leading to the formation of new infectious virus particles (9). Among the viral proteins produced is the Leader (L) protein, a very short protein of 67–76 amino acids, which is cleaved off from the amino-terminal end of the viral polyprotein. Despite its lack of enzymatic activity, the Leader protein is multifunctional as it blocks interferon gene transcription (10, 11), inhibits PKR activation (12, 13) and disrupts nucleocytoplasmic trafficking (NCT)(14–16). In the case of TMEV, these activities were found to depend on L protein ability to interact with cellular partners via two short linear motifs (SLiMs). On the one hand, L interacts with cellular protein kinases of the p90-ribosomal S6 kinase (RSKs) family through a DDVF motif located in its central acidic region (Fig 1). By doing so, L recruits RSKs and maintains these kinases in an activated state (17). L also competes for RSK binding with DDVF containing cellular proteins and thereby dysregulates the RAS-ERK MAP kinase pathway (18). On the other hand, L interacts with the essential components of the nuclear pore complex (NPC) RAE1 and NUP98 (19) through an “M-acidic” SLiM (methionine surrounded by acidic residues) located in the C-terminal region of L (referred to as “Theilo-domain”) (Fig 1A). This SLiM contains a critical methionine residue, which corresponds to methionine 60 in TMEV L (Fig 1). When this key residue is mutated (TMEV-L^M60V^), RSK recruitment remains unaffected, but all L-mediated functions are abolished. Thus, by using a combination of two SLiMs, a DDVF and an M-acidic motif, L retargets part of the cellular RSKs to the NPC where RSKs hyperphosphorylate phenylalanine-glycine nucleoporins (FG-NUPs), thereby triggering NPC opening and allowing free diffusion of nuclear and cytoplasmic proteins (Fig 1A). Although no M-acidic SLiM was delineated in EMCV L, this protein was shown to bear an NPC targeting signal and to trigger RSK-mediated FG-NUPs phosphorylation similarly as TMEV L.

**Fig 1.**
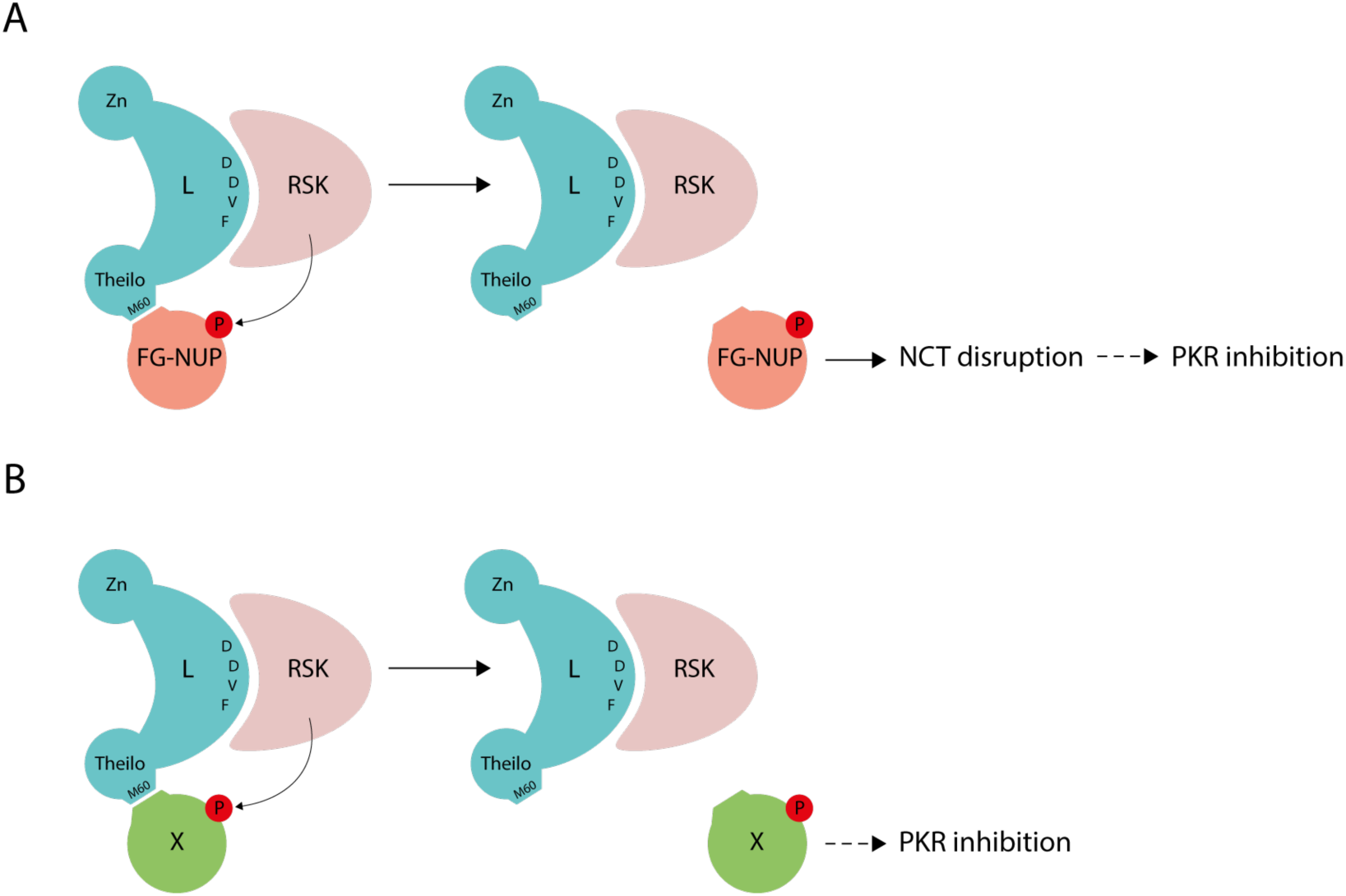
Working models for L-mediated PKR inhibition. A) The Leader protein binds RSK through its DDVF motif and retargets RSK to FG-NUPs through its Theilo-domain, specifically via an “M-acid” SLiM that includes methionine 60 (M60). Hyperphosphorylation of FG-NUPs by RSK ultimately leads to nucleocytoplasmic trafficking (NCT) disruption, and subsequent inhibition of PKR. B) The Leader protein recruits another cellular factor (X) through its Theilo-domain. The phosphorylation of this factor by RSK leads to PKR inhibition. Zn = Zinc finger motif; Theilo = Theilo-domain. P = Phosphorylation.

By triggering the opening of the nuclear pore complex, cardioviruses and other picornaviruses can disrupt the normal trafficking of signaling molecules such as the transcription factors IRF3 and IRF7, thereby effectively impairing the antiviral response (14, 20–22). They can also facilitate the release of nuclear factors such as PTB, PCBP2 and other RNA-binding proteins, which were shown to contribute to viral genome translation and replication in the cytoplasm (14, 22–25).

As both PKR inhibition and NCT disruption by the Leader protein depend on the ability of L to recruit RSKs and on the integrity of the Theilo-domain of L, we asked whether these two activities are connected. PKR inhibition could be a consequence of NCT disruption (Fig 1A) or occur independently of NCT disruption. In the latter case, the Theilo-domain of L would interact with a cellular factor distinct from RAE1-NUP98, whose phosphorylation would lead to PKR inhibition (Fig 1B). As we observed a strong association between PKR inhibition and NCT disruption, we analyzed whether NCT impacted PKR regulation in infected as well as in physiological conditions.

## Results

### Nucleocytoplasmic trafficking disruption is associated with PKR inhibition

To test the extent to which L-induced PKR inhibition is linked to NCT disruption, NCT perturbation was induced in an L-independent manner. Therefore, we used a recombinant virus expressing Coxsackie virus B3 (CBV3) protease 2A (2A^PRO^), which was shown to cleave several FG-NUPs including NUP98, thereby disrupting the integrity of the NPC and impairing nucleocytoplasmic trafficking (26). We used recombinant EMCVs expressing either the wild-type (2A^PRO^) or the catalytically inactive (2A^MUT^) CBV3 2A protease. In these viruses referred to as EMCV-2A^PRO^ and EMCV-2A^MUT^, mutations introduced in the Zinc finger (Zn) and in the DDVF motif (F48A) of L (27) (Fig 2A) prevented L-mediated PKR inhibition and NCT disruption.

**Fig 2.**
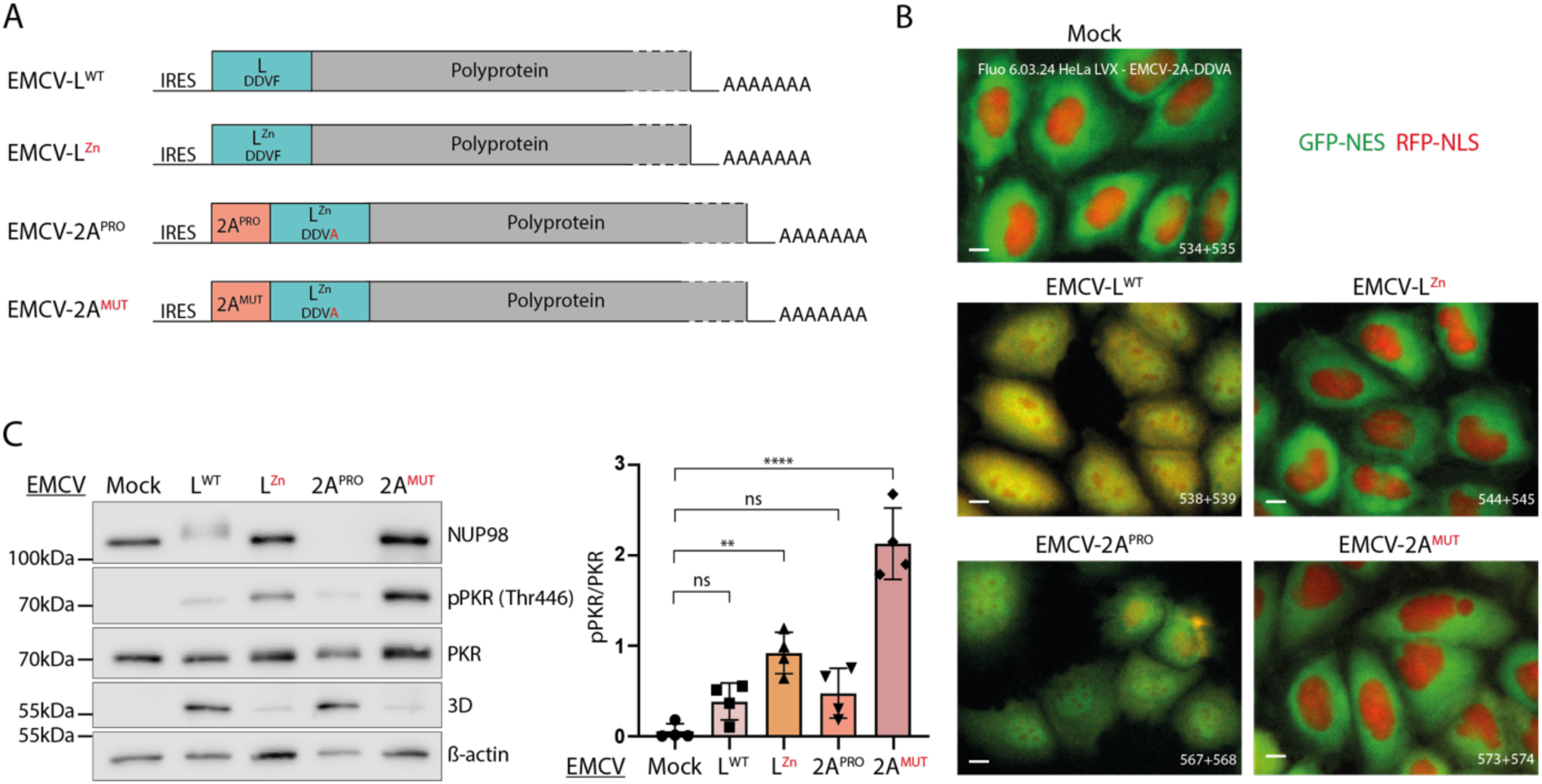
Triggering nucleocytoplasmic trafficking perturbation with an EMCV-2A^PRO^ recombinant virus. A) Schematic representation of EMCV-L^WT^, EMCV-L^Zn^ (controls), EMCV −2A^PRO^, and EMCV-2A^MUT^ genomes. In the EMCV-2A recombinants, a cleavage site for EMCV-3C protease was inserted after the 2A sequence to allow its processing from the viral polyprotein and EMCV L was inactivated by mutating both its zinc finger domain (Zn) and its RSK binding motif (DDVA). B) Confocal microscopy images showing the distribution of RFP-NLS and GFP-NES proteins in live HeLa LVX cells infected for 6h with an MOI of 10 PFU/cell. C) Western blot of HeLa cells infected as in B. 3D polymerase was detected as control of infection and ß-actin as loading control. pPKR (Thr446) and total PKR were quantified from the western blots (mean and SD values; n=4). One-way ANOVA was used to compare all samples with the mock-infected sample. P-values: **: p≤0.01; ***: p≤0.001; ns: non-significant.

To analyze NCT, we used live-cell imaging of infected HeLa LVX cells, which stably express GFP in the cytoplasm (2xGFP-NES) and RFP in the nucleus (2xRFP-NLS). These cells were infected for 6h with EMCV-2A^PRO^ and EMCV-2A^MUT^, and with EMCV-L^WT^ and EMCV-L^Zn^, as controls. As previously shown (15), EMCV-L^WT^ but not EMCV-L^Zn^ triggered extensive NCT disruption, as evidenced by the diffusion of both GFP-NES and RFP-NLS proteins (Fig 2B). Extensive NCT disruption also occurred in cells infected with the recombinant virus expressing active (2A^PRO^) but not catalytically inactive (2A^MUT^) CVB3 protease 2A. These data confirm the ability of CVB3 protease 2A to trigger NCT disruption, when expressed from the recombinant EMCV.

Association between NCT disruption and PKR activation was analyzed by western blot in HeLa cells infected for 6h with those viruses. Consistent with their ability to cause NCT disruption, EMCV-L^WT^ but not EMCV-L^Zn^ triggered NUP98 hyperphosphorylation (migration shift)(Fig 2C) and EMCV-2A^PRO^ but not EMCV-2A^MUT^ triggered a rapid NUP98 cleavage (band disappearance)(Fig 2C). Interestingly, PKR phosphorylation (pPKR) was clearly less intense in cells infected with EMCV-L^WT^ and EMCV-2A^PRO^, than cells infected with EMCV-L^Zn^ and EMCV-2A^MUT^ (Fig 2C), showing that PKR inhibition is associated with NCT disruption.

In the above experiment, it cannot be excluded that PKR inhibition by EMCV-2A^PRO^ might be related to the cleavage of a factor promoting PKR activation, rather than to NCT disruption. We thus designed a second system, unrelated to 2A^PRO^, to trigger NCT disruption and analyze the associated PKR inhibition. To do so, we constructed a TMEV recombinant virus expressing a chimeric protein consisting of the N-terminal part of TMEV-L (L^1-59^), linked to the C-terminal region of SARS-CoV-2 ORF6 (ORF6^48-61^) via a flexible GS linker (Fig 3A). L^1-59^ encompasses the RSK binding motif DDVF but is not sufficient to trigger NCT disruption or PKR inhibition, since it lacks the Theilo-domain. ORF6^48-61^ is known to interact with the RAE1-NUP98 component of the NPC (28) but does not affect PKR activity. The L^1-59^ moiety of the chimera is expected to recruit RSK via the DDVF motif while the ORF6^48-61^ moiety would target the complex toward the nuclear pore complex, where RSK could phosphorylate nucleoporins and trigger NCT disruption. A series of mutants were used as negative controls, including the TMEV-L^M60V^ mutant which is known to suppress the protein’s activities, the complete ORF6 sequence replacing L (TMEV-ORF6), the N-terminal region of L alone (TMEV-L^1-59^), and L-ORF6 constructs with mutations either in the DDVF motif (TMEV-L^F48A^-ORF6) or in the ORF6 methionine 58 (TMEV-L-ORF6^M58A^), which is essential for the interaction with NPC components (28). The last 10 amino acids of L (67–76) containing the 3C cleavage site were inserted at the end of the engineered proteins to promote proper cleavage of the viral polyprotein (Fig 3A).

**Fig 3.**
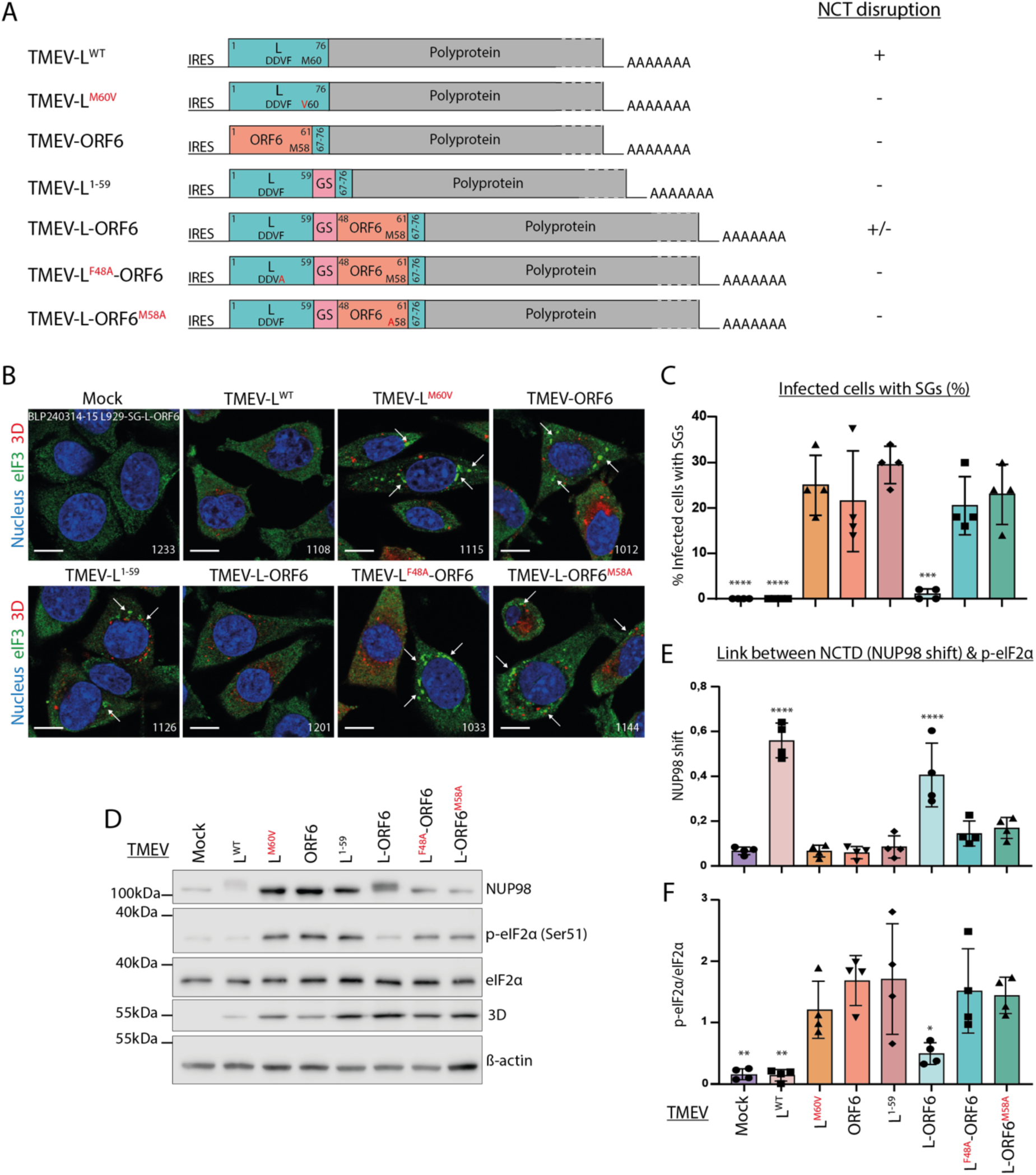
Triggering nucleocytoplasmic trafficking perturbation with a TMEV-L-ORF6 recombinant virus. A) Schematic representation of TMEV recombinant viruses expressing chimeric L-ORF6 proteins and mutants thereof. The impact of the constructs on nucleocytoplasmic trafficking in infected live HeLa LVX is summarized in the column entitled “NCT disruption”, next to the constructs. B) Confocal microscopy images showing SG formation in L929 cells infected for 5h30 with 2 PFU per cell of the indicated viruses. Cells were immunostained for eIF3 (SG marker) and viral polymerase 3D, as a control of infection. Scale bar: 10µm. C) Quantification of cells containing SGs (mean and SD values; n=4). D) Western blot showing the detection of eIF2α phosphorylation in lysates of L929 cells infected as in B. 3D polymerase was detected as control of infection and ß-actin as loading control. E) NUP98 hyperphosphorylation was quantified as the ratio between upward shifted NUP98 and total NUP98 (mean and SD values; n=4). F) p-eIF2α and total eIF2α were quantified (mean and SD values; n=4). One-way ANOVA was used to compare all samples with TMEV-L^1-59^. Shown are significant differences.

NCT disruption by the recombinant viruses was assessed by live-cell imaging of infected HeLa LVX cells (Supplemental Fig S1A). As expected, none of the negative control viruses triggered NCT disruption. In contrast, TMEV-L-ORF6 caused a reproducible NCT disruption though modest compared to that induced by TMEV-L^WT^, characterized by RFP-NLS but not GFP-NES redistribution. Whether partial NCT disruption by TMEV-L-ORF6 is associated with PKR inhibition was assessed in infected murine L929 cells, which are particularly sensitive to infection by the viruses used (29). Since anti-mouse PKR Thr446 phosphospecific antibodies are not available, we used stress granule formation and eIF2α Ser51 phosphorylation as readouts for PKR activation.

SG formation was assessed by staining the eukaryotic translation initiation factor 3 (eIF3) which is a key component of SGs. SG formation was highly decreased in cells infected with TMEV-L^WT^ and TMEV-L-ORF6 viruses compared to cells infected with the control viruses (Fig 3B-C), suggesting that the L-ORF6 chimera inhibited PKR to some extent, in line with its ability to affect NCT. Accordingly, western blot analysis showed less eIF2α phosphorylation (p-eIF2α) in cells infected with TMEV-L^WT^ or TMEV-L-ORF6 than in cells infected with the control viruses (Fig 3D-F). Western Blot analysis also confirmed that both TMEV-L^WT^ and TMEV-L-ORF6, but not the control viruses, induced some extent of NUP98 hyperphosphorylation (migration shift)(Fig 3D-E).

While the effect on NCT disruption was more modest with the TMEV-L-ORF6 recombinant than with EMCV-2A^PRO^, the data nonetheless indicate a consistent association between altered NCT and reduced PKR activation.

To exclude the possibility that PKR inhibition is the cause of NCT perturbation, we analyzed L protein-mediated NCT perturbation in HeLa PKR-KO-LVX cells which express RFP-NLS and GFP-NES but not PKR (Fig 4). GFP-NES and RFP-NLS redistribution was observed in the PKR-KO cells infected with TMEV-L^WT^ but not with TMEV-L^M60V^. These results indicate that NCT disruption is not caused by PKR inhibition and, in contrast, strongly suggest that PKR inhibition is a consequence of NCT disruption.

**Fig 4.**
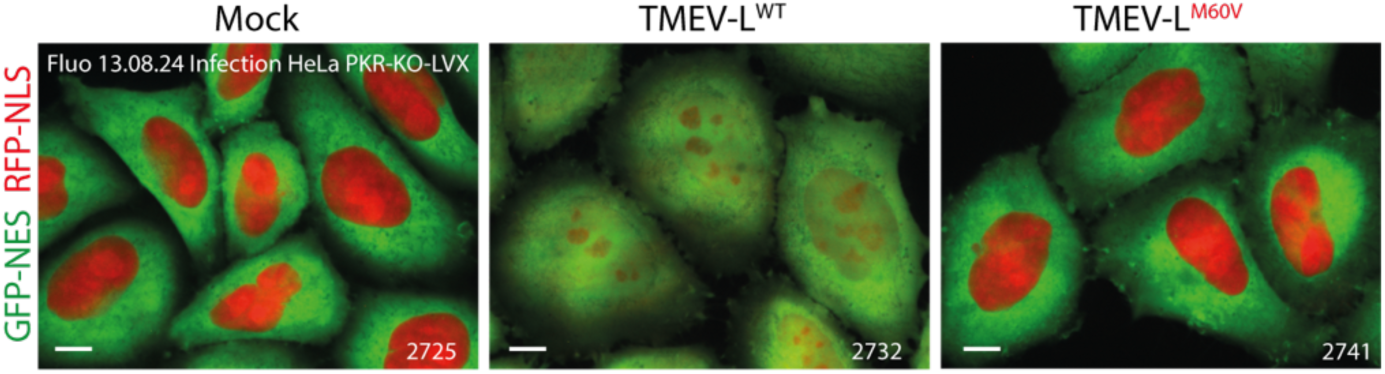
PKR inhibition does not mediate nucleocytoplasmic transport disruption in TMEV-infected cells. Confocal microscopy images showing the distribution of RFP-NLS and GFP-NES in live HeLa PKR-KO LVX cells infected with TMEV-L^WT^ or TMEV-L^M60V^ for 8h at an MOI of 5 PFU/cell. Scale bar: 10µm.

### PKR diffuses into the nucleoli of cells infected with TMEV-L^WT^ and during mitosis

Since PKR is known to be a cytoplasmic protein, its inhibition by the NCT disruption was unexpected. We thus examined the subcellular localization of PKR and of its activated form (pPKR) in infected and non-infected HeLa cells by fluorescence microscopy. On the one hand, PKR was detected using a polyclonal antibody whose specificity was validated using HeLa PKR-KO cells (Fig 5). On the other hand, PKR localization was followed using a split-GFP system where PKR-KO HeLa cells stably expressed the large fragment of GFP (S1-10) as well as PKR fused to the small GFP fragment (S11) (“HeLa PKR-S11” cells) (Fig 5C). In these cells, activated PKR could be detected using an anti-PKR phospho-Thr446 antibody. At validation, the latter antibody showed, however, some (weak) background in the nucleoplasm of PKR-KO cells, but it clearly labeled dotted structures in the cytosol in HeLa M cells infected with TMEV-L^M60V^, known to activate PKR (Fig 5C). In support of the specificity of this antibody, the pPKR signal overlapped almost perfectly (97.7%) that of PKR-S11 (Fig 5C-D).

**Fig 5.**
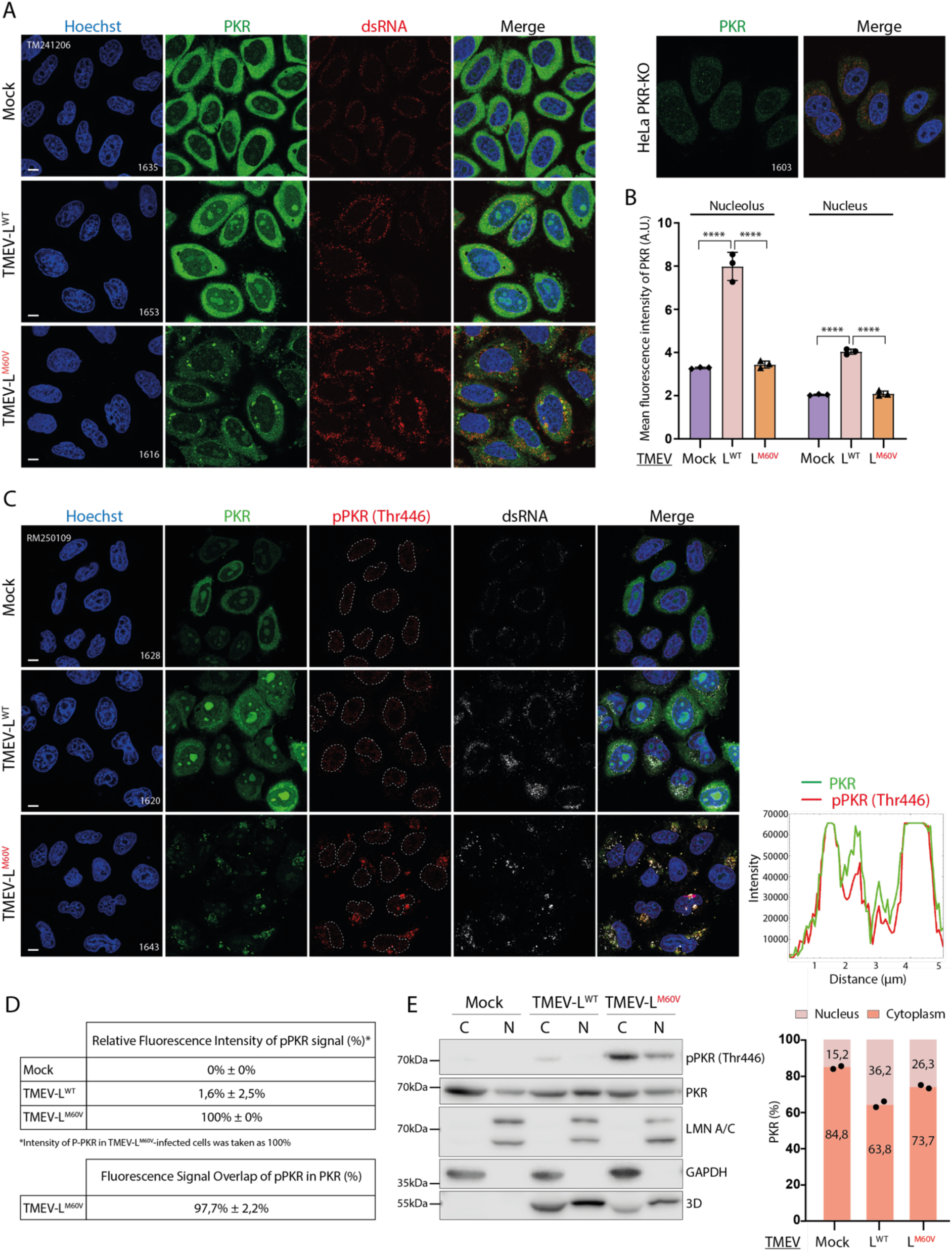
Nucleolar delocalization of PKR during TMEV infection. A) Confocal microscopy images showing PKR subcellular localization in HeLa cells infected for 12h with 5 PFU per cell of TMEV-L^WT^ or -L^M60V^. Cells were immunostained for PKR and dsRNA as control of infection. HeLa PKR-KO cells are shown as a negative control. Scale bar: 10µm. B) Graphs showing the mean fluorescence intensity of PKR in the nuclei and nucleoli (mean and SD values; n=3). One-way ANOVA was used to compare all samples with TMEV-L^WT^. C) Left panel: Confocal microscopy images showing PKR and pPKR subcellular localization in HeLa PKR-S11 cells infected with TMEV-L^WT^ or -L^M60V^ viruses as in A. Cells were immunostained for pPKR and dsRNA, as control of infection. Scale bar: 10µm. Right panel: Intensity vs distance plots of PKR and pPKR in HeLa PKR-S11 cells infected with TMEV-L^M60V^ virus as quantified from the experiment shown in the right panel (pink arrow). D) Table showing the relative fluorescence intensity of pPKR and the percentage of fluorescence signal of pPKR that overlapped with fluorescence signal of PKR. For the relative fluorescence intensity of pPKR, pPKR in TMEV-L^M60V^-infected cells was taken as 100%. E) Western blot analysis of subcellular fractionation of HeLa cells infected with TMEV-L^WT^ or -L^M60V^ viruses for 10h at an MOI of 2.5 PFU/cell. To assess the purity of the fractions obtained, samples were stained for nucleus (Lamin A+C) and cytoplasm (GAPDH) markers. Viral polymerase 3D was detected as control of infection. C = cytoplasm fraction; N = nuclear fraction. PKR signals were quantified from western blot and adjusted according to sample dilution, then expressed as a percentage of total PKR (n=2).

Both native PKR detection (Fig. 5A) and split-GFP detection (Fig. 5C) revealed that, in addition to its mainly cytoplasmic localization, PKR partially maps to nucleoli in uninfected cells. Nucleolar localization of PKR significantly increased upon infection with TMEV-L^WT^ virus but not with TMEV-L^M60V^ which lacks the ability to disrupt NCT (Fig 5A-C). Interestingly, according to phospho-PKR immunostaining, nucleolar PKR was activated neither in non-infected nor in TMEV-L^WT^-infected cells. These results were confirmed by subcellular fractionation experiments showing a higher proportion of PKR in the nuclear fraction of cells infected with TMEV-L^WT^ than in the nuclear fraction of cells infected with TMEV-L^M60V^ or of non-infected cells (Fig. 5E). Again, nuclear PKR was not phosphorylated. Taken together, these results suggest that the disruption of nucleocytoplasmic trafficking triggered by L leads to PKR delocalization to nucleoli, where PKR is not activated.

Interestingly, live imaging of HeLa PKR-S11 cells revealed that, in the absence of infection, PKR accumulates in the nucleus during mitosis and that a substantial portion remains localized within the nucleolus during interphase following mitosis (Fig 6A). In regular HeLa cells, immunostaining showed that PKR is predominantly localized within the nuclear region during the prometaphase and metaphase stages of mitosis (Fig 6B), during which PKR does not appear to be activated (Fig 6C).

**Fig 6.**
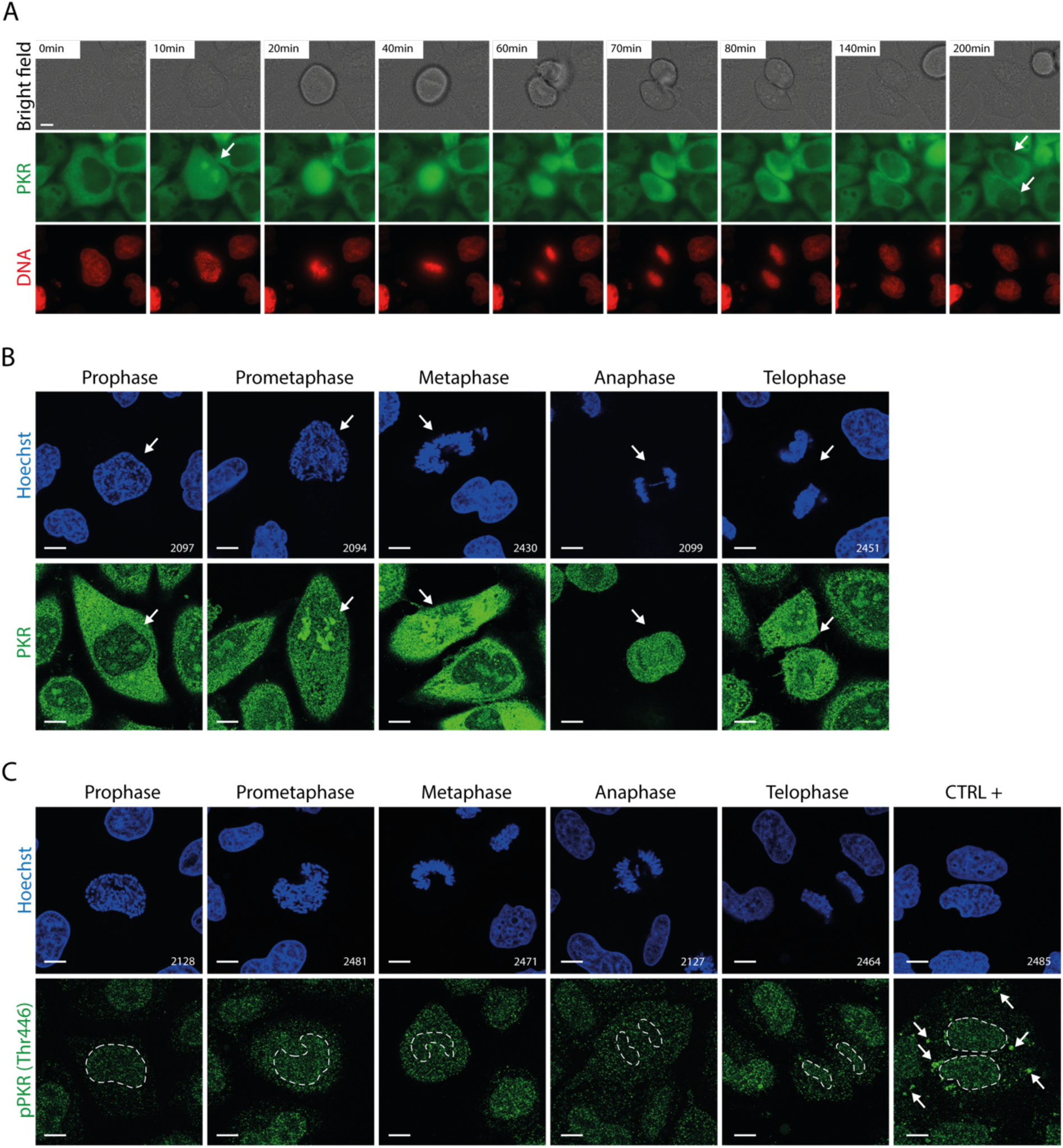
Nucleolar delocalization of PKR during cellular mitosis. A) Time lapse live-cell imaging of HeLa PKR-S11 cells. Arrows point to nuclear/nucleolar PKR condensates. Scale bar: 10µm. B) Confocal microscopy images showing the immunostaining of PKR in HeLa cells during mitosis. Arrows point to a cell undergoing mitosis. C) Confocal microscopy images showing the immunostaining of pPKR in HeLa cells during mitosis. The different mitosis phases were defined based on Hoechst staining. HeLa cells infected for 12h with 5 PFU per cell of TMEV-L^M60V^ were used as positive control. Arrows point to the pPKR signal in the positive control. Scale bar: 10µm.

Taken together, these results suggest that cardioviruses, by triggering the hyperphosphorylation of FG-NUPs, mimic a process taking place during mitosis, which results in the opening of NPCs and the free diffusion of proteins, including PKR, which accumulates to some extent in the nucleoli without being activated.

### Nucleolar localization is driven by the dsRNA binding motifs of PKR

To assess whether nucleolar localization of PKR is associated with dsRNA recognition, we generated HeLa cells transduced to express the dsRNA binding motifs of PKR (N-terminal residues 1-167) fused to 2xmCherry (“HeLa DSRBM PKR-mCherry”). Live-cell imaging of infected HeLa DSRBM PKR-mCherry showed the same observation as with full-length PKR. The mCherry fused to the DSRBM of PKR relocalized to the nucleoli upon infection with TMEV-L^WT^ virus but not with TMEV-L^M60V^ which lacks the ability to disrupt NCT (Fig 7). These observations suggest that the nucleolar relocalization of PKR is associated with recognition of dsRNA or RNA structures mimicking dsRNA within the nucleoli. Additionally, in TMEV-L^M60V^-infected cells, the mCherry-tagged DSRBM of PKR forms distinct perinuclear puncta, which may represent viral replication sites or stress granule–like structures containing dsRNA (Fig 7). These pattern differences suggest that in TMEV-L^WT^ infection, viral RNA is either shielded or rendered inaccessible, supporting a model in which TMEV-L^WT^ employs dsRNA masking to evade host detection.

**Fig 7.**
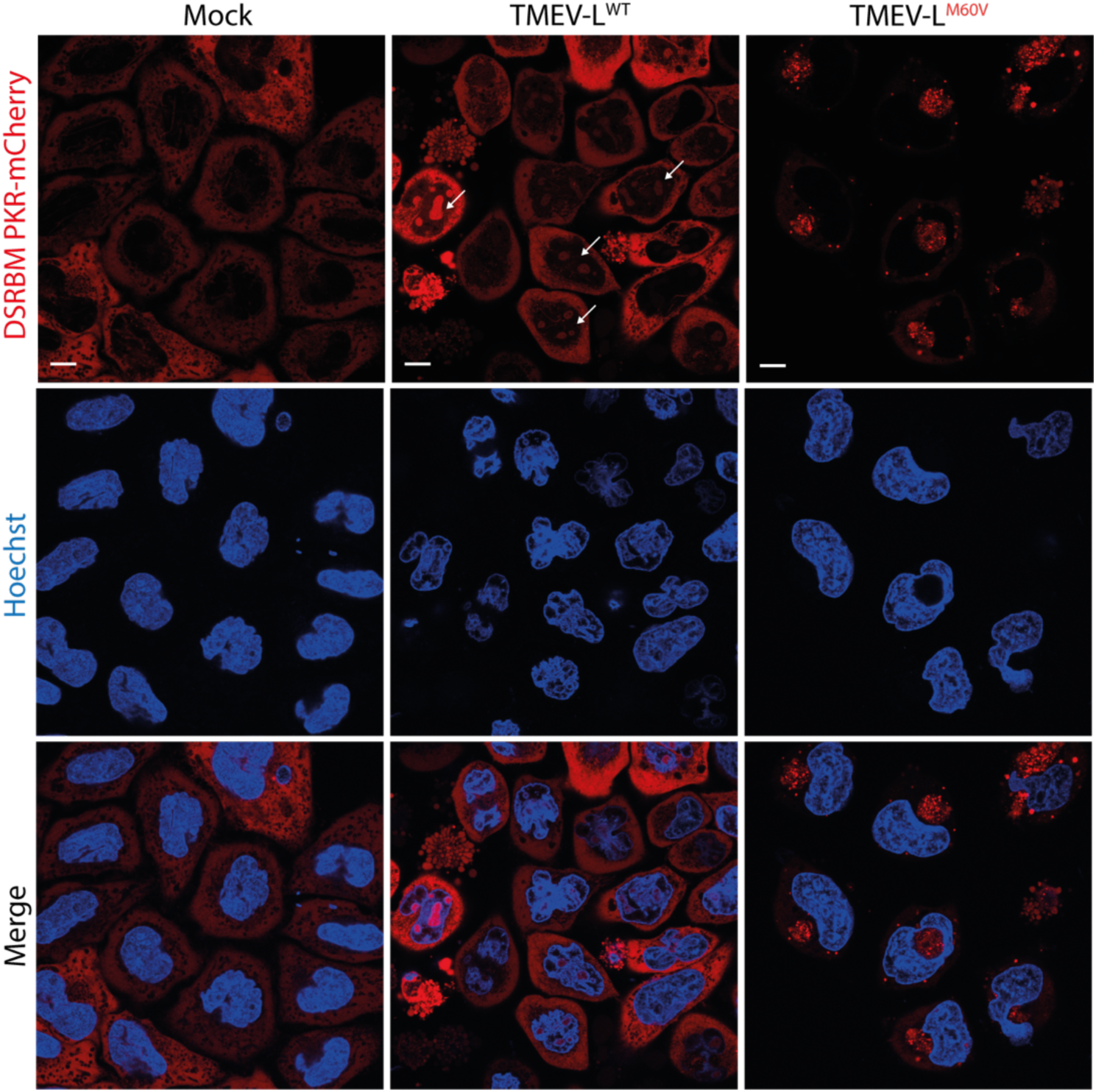
PKR nucleolar localization is driven by its dsRNA binding motifs. Live-cell imaging of HeLa DSRBM PKR-mCherry infected with TMEV-L^WT^ or -L^M60V^ for 12h at an MOI of 5 PFU/cell. White arrows show nucleolar localization of DSRBM PKR-mCherry. Scale bar: 10µm.

### Unclear association between TMEV-L^M60V^-induced SGs and PKR activation

Stress granules were reported to act as a platform for the activation and integration of several antiviral signaling pathways including the PKR-eIF2α pathway (30, 31). We therefore investigated the proximity between phospho-PKR and SGs in HeLa cells infected with TMEV-L^M60V^. As previously shown by Borghese et al. (12), our immunofluorescence analysis showed that in TMEV-L^M60V^-infected cells, PKR and its activated form are localized in two distinct groups of granules, which differ both in their localization and pPKR content (Fig 8A-B): perinuclear granules containing G3BP1, PKR and some amounts of phospho-PKR, and peripheric granules, generally more diffuse, containing PKR, pPKR and small amounts of G3BP1 (Fig 8A-C). Phospho-PKR showed, however, only 55% co-localization with G3BP granules. Based on these results, it remains difficult to conclusively determine whether these stress granule-like structures function as an effective antiviral signaling platform or whether PKR is just trapped in SGs as other RNA-binding proteins.

**Fig 8.**
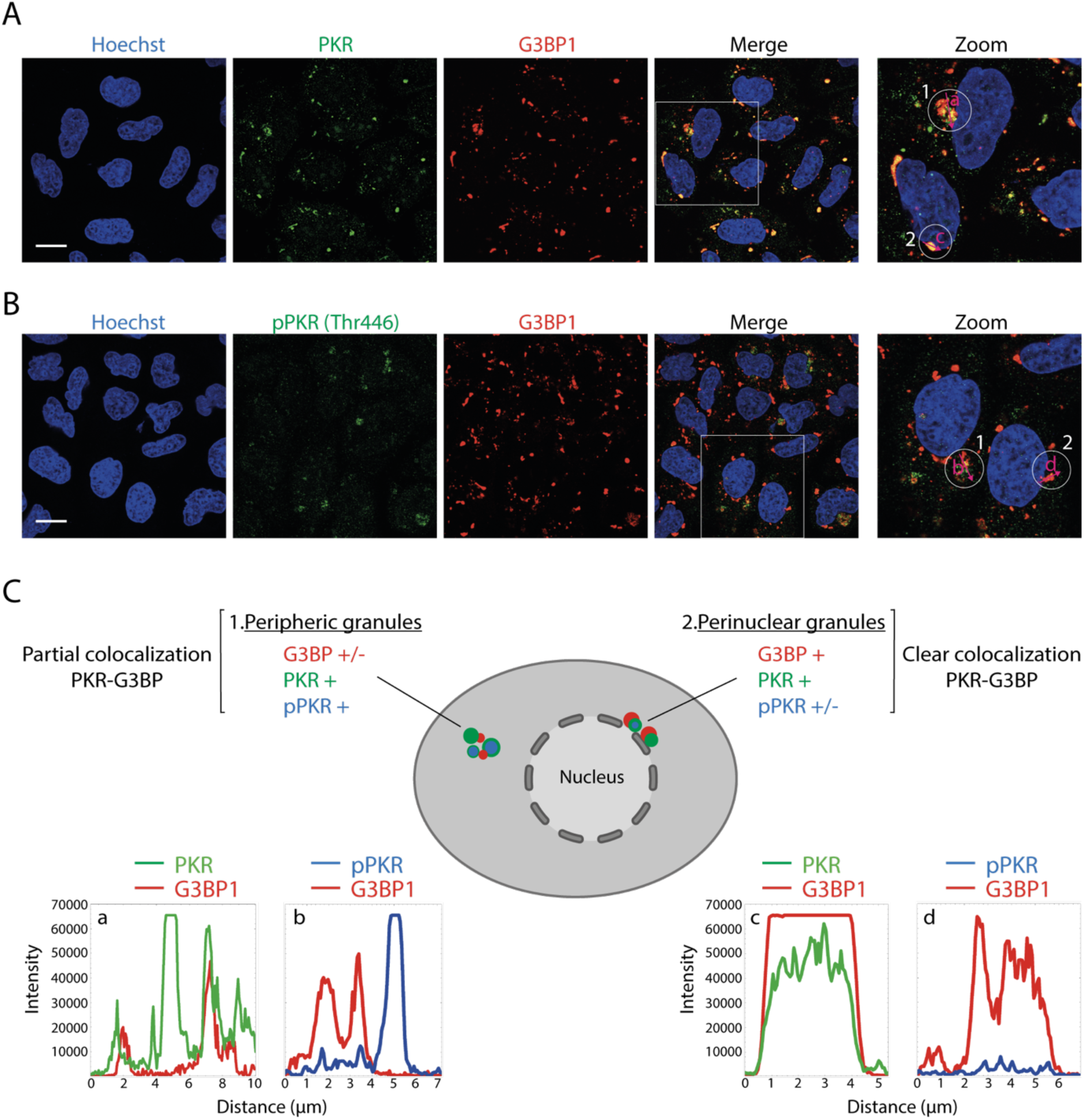
PKR and its activated form are contained into SG-like structures induced by TMEV-L^M60V^ infection. A) Confocal microscopy images showing the co-immunostaining of PKR and G3BP1 in HeLa cells infected for 12h with 5 PFU/cell of TMEV-L^M60V^. G3BP1 was detected as a canonical marker of SG. Scale bar: 10µm. B) Confocal microscopy images showing the co-immunostaining of pPKR and G3BP1 in HeLa cells infected as in A. Scale bar: 10µm. C) Cartoon summarizing the observations with intensity vs distance plots of PKR (green) and G3BP1 (red) or pPKR (blue) and G3BP in HeLa cells infected as quantified from the experiments shown in A and B (pink arrow).

### Association between PKR inhibition and nuclear RBPs diffusion into the cytoplasm

It is well established that cardioviruses, by inducing the opening of the NPC, promote the cytoplasmic translocation of several nuclear RNA-binding proteins, including Pyrimidine tract-protein 1 (PTB) and Src-associated in mitosis of 68 kD (Sam68). These proteins, once relocated to the cytoplasm, are thought to support essential viral processes such as viral RNA translation and genome replication (14, 23, 32, 33). Extending these findings, we recently observed that two additional nuclear proteins, NONO and SFPQ – both paraspeckle-associated proteins containing RNA recognition motifs – similarly diffuse to the cytoplasm during TMEV infection (Supplemental Fig S2A-B). This growing list of mislocalized nuclear proteins during infection suggests a broader viral strategy to manipulate host cell function by altering the subcellular localization of important proteins. To identify proteins acting as potential PKR competitors/inhibitors in cells infected with TMEV-L^WT^, we generated HeLa cells (“HeLa PKR-APEX”) expressing a fusion between PKR and an engineered ascorbate peroxidase (APEX), which enables promiscuous biotinylation of proximal proteins (Fig 9A) (34).

**Fig 9.**
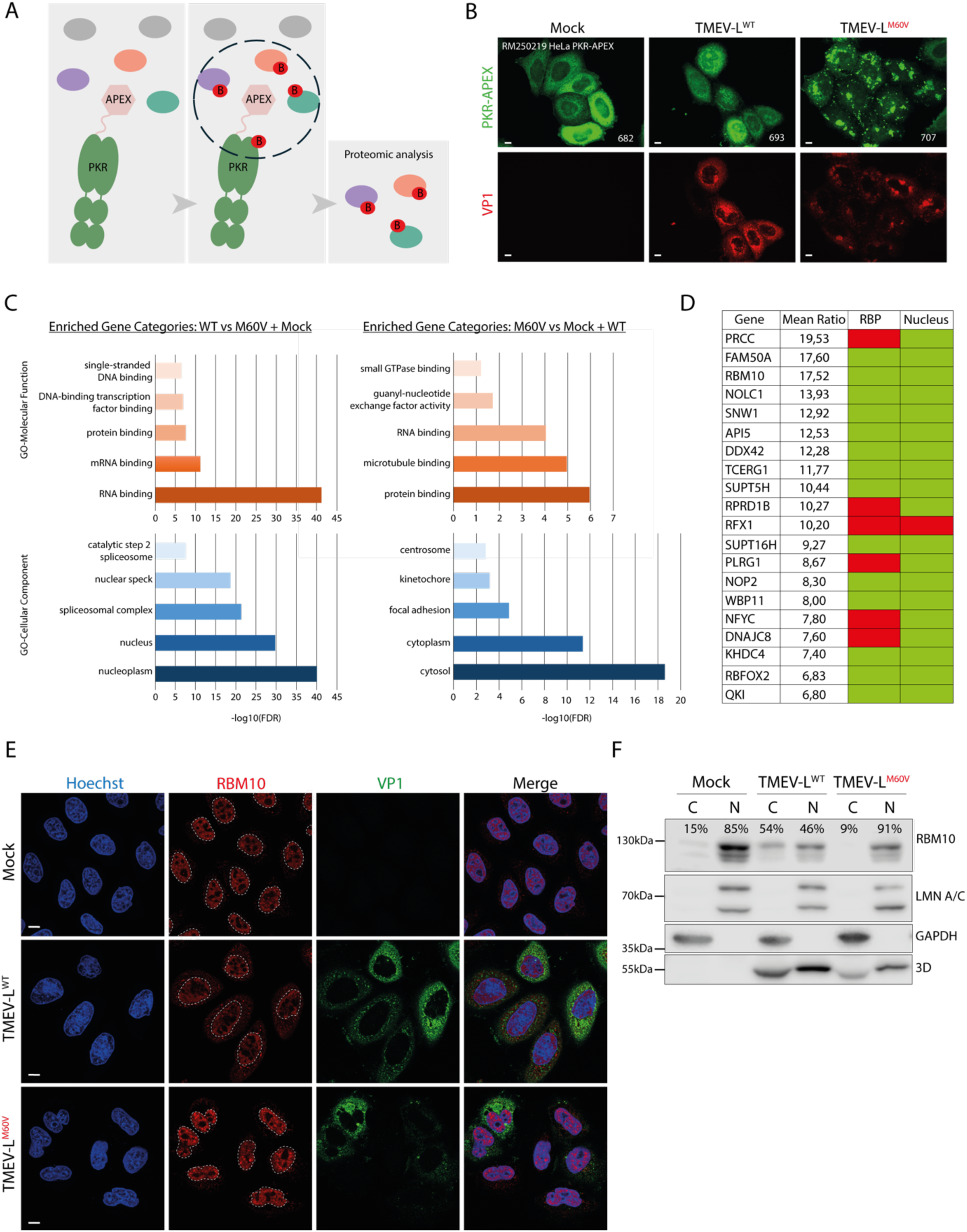
Cytoplasmic diffusion of nuclear RNA-binding proteins during TMEV infection. A) Cartoon showing PKR-APEX-mediated proximity biotinylation of proteins. Addition of biotin-phenol for 1h and hydrogen peroxide for 2 minutes to HeLa cells results in biotinylation of proteins within closed proximity of PKR-APEX. Biotinylated proteins can then be purified by streptavidin pulldown and identified by mass spectrometry. B) Confocal microscopy images showing the co-immunostaining of PKR and VP1. VP1 was detected as a control of infection. Scale bar: 10µm. C) Gene Ontology (GO) enrichment analysis as a bar graph, with the x-axis representing -log₁₀(FDR) (False Discovery Rate) and the y-axis displaying GO molecular function or GO cellular component terms. The -log₁₀(FDR) transformation enhances the visualization of statistical significance, where higher values indicate stronger enrichment. D) Table summarizes the top 20 candidates, highlighting their RNA-binding capacity and nuclear localization. Green indicates confirmed nuclear localization or RNA-binding capacity; red indicates absence of nuclear localization or RNA-binding ability. E) Confocal microscopy images showing the co-immunostaining of RBM10 and VP1 in HeLa cells infected with TMEV-L^WT^ or -L^M60V^ viruses for 12h at an MOI of 5 PFU/cell. VP1 was detected as a control of infection. Scale bar: 10µm. F) Western blot analysis of subcellular fractionation of HeLa cells infected with TMEV-L^WT^ or -L^M60V^ viruses for 10h at an MOI of 2.5 PFU/cell. To assess the purity of the fractions obtained, samples were stained for nucleus (Lamin A+C) and cytoplasm (GAPDH) markers. Viral polymerase 3D was detected as control of infection. C = cytoplasm fraction; N = nuclear fraction. RBM10 signals were quantified and adjusted according to sample dilution, then expressed as a percentage of total RBM10.

After testing that the PKR-APEX construct exhibited expected regulation – being inhibited in TMEV-L^WT^-infected cells and activated in TMEV-L^M60V^-infected cells (Supplemental Fig S3A) – and was localized as wild-type PKR (Fig 9B), we performed an APEX-mediated biotinylation experiments in TMEV-infected cells. To do so, HeLa PKR-APEX cells were mock-infected or infected with 5 PFU per cell of TMEV-L^WT^ or -L^M60V^ viruses for 4, 6, 8 or 10 hours in presence of biotin-phenol. After hydrogen peroxide activation of APEX, biotinylated proteins were captured using streptavidin beads, and western blot analysis confirmed their capture in the pulled-down fraction (Supplemental Fig S3B). The proteins were then identified by mass spectrometry (Pride repository dataset identifier PXD066206 and 10.6019/PXD066206) and sorted according to their peptide spectrum match (PSM) number. Protein abundance in TMEV-L^WT^–infected cells was quantified relative to both uninfected and TMEV-L^M60V^-infected cells (for more detail on the calculations see material and methods) at each time point. These relative abundance ratios were averaged across all time points, and proteins were ranked based on their mean ratios. The top 150 candidates were subsequently selected for Gene Ontology (GO) enrichment analysis. Figure 9C presents the results of GO enrichment analysis as a bar graph, where higher values indicate stronger enrichment. Our analysis revealed significant enrichment of proteins with RNA-binding capacity as well as proteins localized in the nucleus. These results suggest that nuclear and RNA-binding proteins are more abundant in the vicinity of PKR in TMEV-L^WT^-infected cells than in mock- and TMEV-L^M60V^-infected cells. This observation may be explained by PKR relocalization to the nucleolus, by diffusion of these proteins into the cytoplasm, or by a combination of both. To further investigate these findings, we selected two of the highest-scoring candidates (Fig 9D), RBM10 and PRCC, and performed either immunofluorescence (Fig 9E; Supplemental Fig S3C) or subcellular fractionation (Fig 9F) analysis on HeLa cells infected with TMEV-L^WT^ or TMEV-L^M60V^ viruses. The results confirmed the cytoplasmic diffusion of RBM10 and PRCC in TMEV-L^WT^-infected cells, in contrast to their nuclear localization in mock- and TMEV-L^M60V^-infected cells. These results demonstrated that L-mediated NCT disruption results in the release of nuclear RNA-binding proteins, providing further insight into the mechanisms by which cardioviruses evade host antiviral responses.

Taken together, our findings strongly support the hypothesis that TMEV-induced NCT disruption contributes to PKR inhibition. As nucleocytoplasmic trafficking is impaired, PKR appears to be partially recruited to the nucleolus, where it remains inactive. Additionally, L-induced NCT promotes the extensive release of nuclear RNA-binding proteins into the cytoplasm. These proteins may either compete with PKR for viral dsRNA binding, modify viral RNA to prevent PKR recognition, or directly interact with and modulate PKR to block its activation (Fig 10). In stark contrast, infection with the TMEV-L^M60V^ virus, which fails to disrupt nucleocytoplasmic trafficking, maintains the proper nuclear localization of RNA-binding proteins, ensuring that PKR remains in the cytoplasm, where it can effectively detect and bind to double-stranded RNA, thus becoming activated. These results provide compelling evidence for the intricate interplay between viral infection, nucleocytoplasmic trafficking, and the regulation of the PKR pathway.

**Fig 10.**
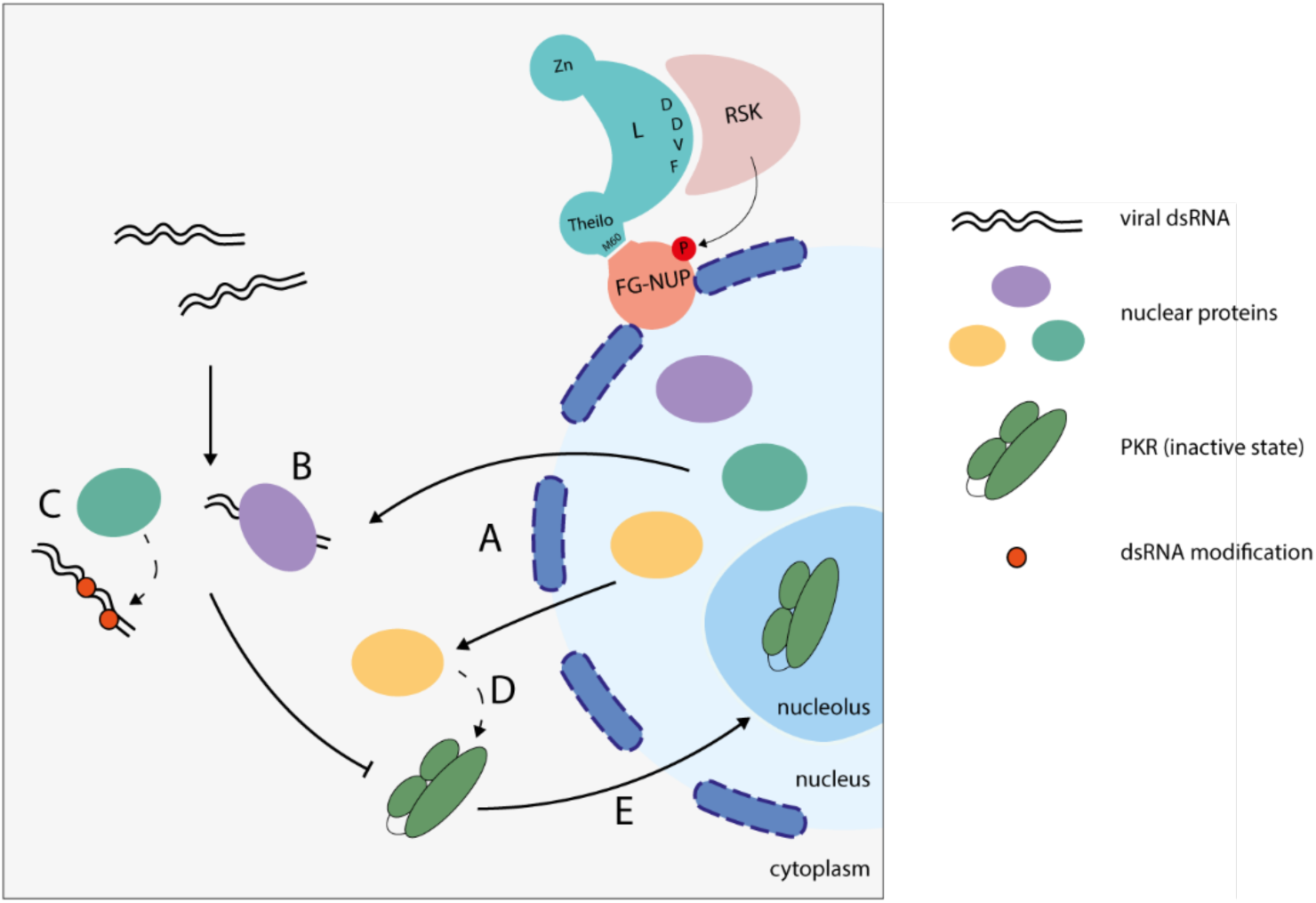
NCT at the origin of PKR inhibition. Nuclear protein diffusion into the cytoplasm during infection (A) may sequester viral dsRNA (B), modify viral dsRNA (C), or interact with and modify PKR (D), thereby preventing PKR activation. Moreover, impaired nucleocytoplasmic trafficking results in partial localization of PKR to the nucleolus, where it remains inactive (E).

## Discussion

In this study, we investigated the mechanisms underlying PKR inhibition during Theiler’s murine encephalomyelitis virus infection, focusing on the role of the viral L protein in modulating nucleocytoplasmic trafficking. Our findings demonstrate that this inhibition is closely linked to the disruption of nucleocytoplasmic trafficking orchestrated by the viral L protein (15). Using recombinant viruses engineered to trigger alternative trafficking defects (27), we show that PKR inhibition is likely a direct consequence of L-induced nucleocytoplasmic trafficking disruption.

Although PKR is generally considered to localize and function in the cytoplasm, a distribution supported by the presence of a nuclear export signal (NES) in its N-terminal region (35), we observed that the disruption of nucleocytoplasmic transport leads to its partial relocalization to nucleoli. This observation is consistent with earlier findings by Jeffrey et al. (36) who reported nucleolar localization of PKR in human Daudi cells and in stably transfected mouse NIH 3T3 cells expressing human PKR. Furthermore, Hakki et al. (37) demonstrated that the TRS1 and IRS1 gene products of human cytomegalovirus (HCMV) directly interact with PKR and inhibit its activation by sequestering it in the nucleus, away from both its activator, cytoplasmic dsRNA, and its substrate, eIF2α.

Interestingly, we also found that PKR accumulates in the nuclear region during mitosis, particularly at prometaphase and metaphase. Our data show that nucleolar accumulation of PKR is driven by the RNA-binding domain of the protein, suggesting that PKR binds to nucleolar RNA that forms dsRNA or local dsRNA-like structures. However, despite the high concentration of RNA in nucleoli, we did not detect PKR activation at these stages in contrast to the findings of Kim et al. (38) who reported that, upon nuclear envelope breakdown during mitosis, PKR is activated through binding to dsRNAs formed by inverted Alu repeats.

A possible explanation for the lack of PKR activation in the nucleoli would be that PKR monomers, individually bound to RNA, are hindered from dimerizing – an essential step for autophosphorylation and activation. This model is supported by the findings of Lemaire et al. (39), who demonstrated that an excess of dsRNA inhibits PKR by dispersing it among numerous dsRNA molecules, thereby preventing autophosphorylation between PKR molecules. This sequestration within the nucleolus would therefore constitute a novel viral strategy to spatially inhibit PKR function despite its RNA-rich environment.

An alternative explanation for the absence of PKR activation in the nucleolus could be the presence of RNA molecules that inhibit PKR. Indeed, several RNAs from both viral and endogenous sources have been identified as inhibitors of PKR. Viral RNAs, such as adenovirus VA-I and Epstein-Barr virus EBER-1, prevent PKR activation by binding to its double-stranded RNA-binding motifs, thereby blocking autophosphorylation and downstream signaling – either by sequestering PKR in a monomeric state (VA-I) or by inducing an inactive dimeric form (EBER-1) (40–42). Additionally, the host-derived noncoding RNA nc886 regulates basal PKR activity, existing in two structural conformers with differing inhibitory capacities. Conformer 1 binds strongly to and inhibits PKR, whereas conformer 2 exhibits weak binding and functions as a pseudoactivator (43, 44).

Biomolecular condensates are known to form in response to various cellular stresses, including viral infections. These condensates, and particularly stress granules, have been proposed to act as platforms for orchestrating antiviral innate immune signaling pathways, including those involving PKR (30, 31, 45, 46). Notably, it has been demonstrated that G3BP1, a key nucleating factor of SGs, can induce the formation of SGs in the absence of cellular stress. This spontaneous formation can activate PKR, thereby inducing eIF2*α* phosphorylation and subsequent translational repression (47). These finding establish a link between the mere assembly of stress granules and the activation of PKR. On one hand, SG-like structures might facilitate PKR activation by concentrating RNAs and essential cofactors. On the other hand, they could serve as compartments for sequestration, limiting access to activating viral RNAs. Interestingly, we observed that both total and phosphorylated PKR only partially colocalize with G3BP1. Therefore, it remains unclear whether SGs actively regulate PKR activity or if their formation is a downstream consequence of virus-induced cellular stress. Further studies are needed to elucidate the precise role of SGs in the regulation of PKR and other components of the antiviral immune response.

Additional mechanisms for PKR inhibition are suggested by proximity labeling experiments, which showed that L-mediated disruption of nucleocytoplasmic trafficking leads to the mislocalization of numerous nuclear RNA-binding proteins into the cytoplasm. These RBPs, once relocalized, could compete with PKR for double-stranded RNA molecule detection, thereby limiting its activation. One potential mechanism is that nuclear RBPs bind directly to viral RNAs, shielding them from PKR recognition. Alternatively, specific nuclear enzymes – such as RNA editases, methyltransferases, or other RNA-modifying proteins – may modify viral RNA, making it undetectable by PKR. A third possibility is that certain nuclear proteins may act on PKR itself, either through binding or post-translational modifications, to prevent its activation.

Taken together, our findings reveal a sophisticated, multi-layered viral strategy to evade PKR-mediated antiviral responses. We found that TMEV infection promotes the relocalization of nuclear RNA-binding proteins to the cytoplasm, where they likely compete with PKR for double-stranded RNA binding, thereby preventing its activation. Furthermore, during both TMEV infection and mitosis, PKR accumulates in the nucleoli, where it likely interacts with structured RNAs without becoming activated. These results collectively underscore nucleocytoplasmic trafficking as a critical regulatory mechanism controlling PKR activation. Beyond the specific activity of the Cardiovirus L protein, our study points to a broader principle in which interference with host nucleocytoplasmic transport can significantly alter the subcellular localization and function of immune effectors, such as PKR.

## Material and methods

### Cells

Cells referred to as HeLa cells in this work are HeLa M cells, a subclone of HeLa cells kindly provided by R. H. Silverman (48). HeLa PKR-KO cells, generated by CRISPR-Cas9 genome editing, were described previously (Cesaro et al. 2021). HeLa-LVX and HeLa PKR-KO-LVX cells expressing RFP-NLS and GFP-NES were obtained as described previously (15).

HeLa PKR-S11 cells were obtained by transduction of HeLa PKR-KO cells with the lentivirus BLP2 coding for the strands S1-10 of GFP. After transduction, a cellular clone was selected, which showed low fluorescence in the absence of GFP segment S11, but readily detectable fluorescence after infection with an S11 expressing virus. This cloned was then transduced with the lentivirus FW28 coding for the human PKR fused to the GFP segment S11. After transduction, a cellular clone was selected, which showed easily detectable fluorescence in a pattern that matched typical “PKR granules” found in TMEV-L^M60V^-infected cells.

HeLa PKR-APEX cells were obtained by transduction of HeLa PKR-KO cells with the lentivirus RM15 (see below) coding for the human PKR fused to APEX3. After transduction, a cellular clone was selected, which showed a PKR expression level similar to that of PKR in HeLa cells.

L929 cells were obtained from ECACC (ref 85011425); BHK-21 cells were obtained from ATCC (CCL-10). 293T cells (49) used in this work for lentivirus production, were kindly given by F. Tangy (Pasteur Institute, Paris). HeLa, L929 and 293T cells were maintained in Dubbelco’s Modified Eagle Medium (DMEM high-glucose - Biowest) supplemented with 10% fetal calf serum (FBS, Hyclone-Cytiva), penicillin (100U/ml) and streptomycin (100μg/ml) (Gibco). BHK-21 cells used for virus production were maintained in Glasgow’s Modified Eagle Medium (GMEM, Gibco) with 2.6g/L of tryptose phosphate broth (Gibco), 10% newborn calf serum (Gibco) and penicillin (100U/ml)/streptomycin (100μg/ml). All cell lines were cultured at 37°C, 5% CO2.

### Lentiviral vectors and cell transduction

Expression plasmids and retro/lentiviral expression vectors are presented in Table 1. Retroviral expression vectors were derived from pQCXIH (Clontech) and lentiviral vectors were derived from pCCLsin.PPT.hPGK.GFP.pre. (50).

For lentivirus production, 293T cells were seeded in a 6-cm Petri dish. At ∼80% confluency cells were co-transfected with 2.5µg of lentiviral vector (pBLP2, pFW28 or pRM15), 1.250µg pMDLg/RRE (Gag-Pol), 0.75µg of pMD2-VSV-G, and 0.625µg of pRSV-Rev using TransIT-LT1 (Mirus Bio). 24h and 48h post transfection, supernatant was collected and filtered (0.45µm).

For transduction, cells were typically seeded in a 24-well plate as of 5,000 cells/well and were transduced from 2 to 10 times with 100µL of filtered lentivirus (BLP2, FW28 or RM15).

### Virus production and titration

TMEV viruses used in this work are derivatives of KJ6, a variant of the persistent DA strain (DA1 molecular clone) adapted to grow on L929 cells (29). FB09 carries the M60V mutation in L (L^M60V^), which was shown to abrogate all known L activities (13, 17, 51).

EMCV derivatives (Mengo virus) were obtained from pMC24 (originally named pMC16.1) a cDNA clone kindly provided by Ann Palmenberg (52).

TMEV and EMCV derivatives were produced by reverse genetics from plasmids containing their full-length viral cDNA sequences. To this end, BHK-21 cells were electroporated with viral RNA (*in vitro* transcribed – RiboMax Promega P1300) using a Gene pulser apparatus (Bio-Rad) (1500V, 25µFd, no shunt resistance). Supernatants were collected when cytopathic effect was complete (∼48h post-electroporation). Two to 3 freeze-thaw cycles were made to increase viral release from cells before clarifying the supernatants by centrifugation (20min, 1258g). Viruses were stored at −80°C and titrated by plaque assay in BHK-21 cells as described (4).

Viruses and plasmids used in this study are listed in supplementary table S1.

### Western blot

Proteins in Laemmli buffer were heated at 100°C for 5 min. Protein samples were then run in 10% glycine SDS polyacrylamide gels. Proteins were then transferred to PVDF membranes. For p-eIF2*α* detection, membranes were treated with a signal enhancer (ref 46640 - ThermoScientific) prior to chemiluminescence detection. Membranes were blocked with TBS-5% milk (Regilait) for 1h at RT. Membranes were then incubated over-night with primary antibodies at proper dilution in TBS-5% milk. Primary antibodies used: anti-pPKR (ab32036 Abcam, rabbit, 1/2000), anti-PKR (18-244-1-AP Proteintech, rabbit, 1/4000), anti-NUP98 (N1038 Sigma, rat, 1/1000), anti-ß-actin (A5441 Sigma, mouse, 1/10000), anti-p-eIF2*α* (CST9721 Cell Signaling, rabbit, 1/1000), anti-eIF2*α* (CST9722 Cell Signaling, rabbit, 1/1000), anti-GAPDH (MAB374 Millipore, mouse, 1/10000), anti-Lamin A/C (GTX101127 GeneTex, rabbit, 1/4000) and anti-3D-viral polymerase polyclonal rabbit antibody (See below - 1/2000). Membranes were washed 3 times with TBS-0.1% Tween 20 for 15min before incubation with secondary antibodies in TBS-5% milk for 1h at RT. Secondary antibodies used: HRP-conjugated anti-rabbit (Dako P0448 - 1/5000), HRP-conjugated anti-mouse (Dako P0447 - 1/5000), and HRP-conjugated anti-rat (CST 7077 - 1/5000). Membranes were washed 3 times with TBS-0.1% Tween 20, then once with TBS and revealed with SuperSignal West chemiluminescence substrate (Pico or Dura, ThermoScientific) or with Westar Supernova (Cyanagen). Images were taken with a cooled CCD camera (Odyssey FC-LiCor).

### Subcellular fractionation

A total of 2 × 10^5^ HeLa cells were plated in 5cm diameter culture dishes. Two days after, cells were either mock-infected or infected with TMEV-L^WT^ or TMEV-L^M60V^ for 10 h with an MOI of 2.5. Cytoplasmic and nuclear extracts were prepared according to the instructions of the NE-PER^TM^ nuclear and cytoplasmic extraction kit (ThermoScientific ref 78833).

### Immunostaining

Cells seeded in a 96-well plate (Greiner, 655866 Screenstar) were fixed with PBS-4% PFA for 5min at RT before being permeabilized with PBS-0.2% Triton (ICN Biomedicals Inc.) for 5min. Cells were blocked with TNB blocking reagent (Perkin Elmer) for 1h at RT. Cells were then washed three times with PBS-0.1% Tween 20 before being incubated with primary antibody diluted in TNB for 1h at RT. Primary antibodies: anti-pPKR (MA5-46897 (HL1439)

Invitrogen, rabbit, 1/50), anti-PKR (18-244-1-AP Proteintech, rabbit, 1/800), anti-eIF3 (sc-137214 Santa cruz, mouse, 1/800), anti-dsRNA (mAb K1 EngScicons (Hungary), mouse, 1/400), anti-G3BP1 (611126 BD Biosciences, mouse,1/400), anti-VP1 (F12B3 clone, mouse, 1/25; a kind gift of M. Brahic), anti-PRCC (PA5-53998 Invitrogen, rabbit, 1/50), anti-RBM10 (14423-1-AP Proteintech, rabbit, 1/50) and anti-3D-viral polymerase polyclonal rabbit antibody (See below - 1/800). Cells were then washed three times with PBS-0.1% Tween 20 before being incubated with secondary antibody diluted in TNB for 1h at RT. Secondary antibodies: anti-rabbit Alexa Fluor 488 (A11008 – goat,1/800), anti-rabbit Alexa Fluor 594 (A11037 – goat, 1/800), anti-mouse Alexa Fluor 488 (A11029 – goat, 1/800), anti-mouse Alexa Fluor 594 (A11032 – goat, 1/800) and anti-mouse Alexa Fluor-647 (A32728 – goat, 1/800). Cells were then washed three times with PBS-0.1% Tween 20 and kept in PBS-0.02% sodium azide.

### APEX-mediated biotinylation experiment

A total of 1.5 × 10^6^ HeLa PKR-APEX cells were plated in 10cm diameter culture dishes. Cells were either mock-infected or infected with TMEV-L^WT^ or TMEV-L^M60V^ for 4, 6, 8 or 10 h with an MOI of 5. 1 h before the end of the infection, medium was replaced by fresh medium containing 5mM of biotin-phenol and incubating at 37°C under 5% CO2 for 1 h. Medium was finally replaced by fresh medium containing 1mM of H2O2 for 2min. Cells were then washed two times with 5mL of quenching solution (10mM sodium ascorbate, 10mM sodium azide and 5mM Trolox in PBS) and one time with PBS.

### Streptavidin pulldown

Cells were lysed with 800µL/dish of stringent lysis buffer (50mM Tris-HCl pH 7.6, 500mM NaCl, 0.4% SDS, 1mM DTT, 1 tablet of Pierce phosphatase/protease inhibitor (Thermo Scientific) per 10ml of lysis buffer) for 15 min at room temperature (RT). Lysates were then homogenized by 10 passages through 21G needles. Lysates were then cleared by centrifugation at 12,000g for 10 min at RT. Supernatants were then passed through ZebaSpin columns, as described by the manufacturer (ThermoScientific ZebaTMSpin Desalting Columns 7K MWCO ref 89892), to eliminate free biotin. 25μL (per condition) of protein A/G magnetic beads (Pierce) were added to remove non-specific binding and incubated for 30 min at RT. Supernatants were then transferred to a new tube and a sample of 50μL per condition was mixed with 25μL of 3x Laemmli buffer (cell lysate control). The rest of the supernatant was incubated for 2h at RT with 50μL (per condition) of Streptavidin magnetic beads (Pierce). Streptavidin beads were then washed twice with 2% SDS, once with stringent lysis buffer and once with “normal lysis buffer” (50mM Tris-HCl pH 7.5, 100mM NaCl, 2mM EDTA, 0.5% NP40, 1 tablet of phosphatase/protease inhibitor (Thermo Scientific) per 10ml of lysis buffer).

Beads were then resuspended in 500µL of wash buffer (Tris-HCl 50mM pH7.4, NaCl 100mM, 1 tablet of phosphatase/protease inhibitor (Thermo Scientific) per 10ml of lysis buffer) and heated for 3min at 100°C to allow non-biotinylated protein’s separation from the beads. Beads were finally resuspended in 50µL of 1x Laemmli buffer and heated for 5min at 100°C to allow biotinylated protein’s separation from the beads. Supernatants were then conserved at −20°C.

### Mass spectrometry (Orbitrap Lumos)

Streptavidin pulldown samples were resolved using a 10% Tris-Glycine SDS gel run until 6mm migration in the separating gel. Proteins were colored using PageBlue (Thermo Scientific, 24620). The 6mm bands containing whole proteins were cut into 3 different slices and trypsin digested. Peptides were extracted with 0.1% TFA in 65% ACN and dried in a speedvac.

Peptides were dissolved in solvent A (0.1% TFA in 2% ACN), directly loaded onto reversed-phase pre-column (Acclaim PepMap 100, Thermo Scientific) and eluted in backflush mode. Peptide separation was performed using a reversed-phase analytical column (Acclaim PepMap RSLC, 0.075 x 250 mm,Thermo Scientific) with a linear gradient of 4%-27.5% solvent B (0.1% FA in 98% ACN) for 40 min, 27.5%-50% solvent B for 20 min, 50%-95% solvent B for 10 min and holding at 95% for the last 10 min at a constant flow rate of 300 nl/min on an Vanquish Neo system. The peptides were analyzed by an Orbitrap Fusion Lumos tribrid or Exploris240 mass spectrometer (ThermoFisher Scientific). The peptides were subjected to NSI source followed by tandem mass spectrometry (MS/MS) in Fusion Lumos or Exploris240 coupled online to the nano-LC. Intact peptides were detected in the Orbitrap at a resolution of 120,000. Peptides were selected for MS/MS using HCD setting at 30, ion fragments were detected in the Orbitrap at a resolution of 30,000 (Exeploris240) or in the linear ion trap (LUMOS). A data-dependent procedure that alternated between one MS scan followed by MS/MS scans was applied for 3 seconds for ions above a threshold ion count of 2.0E4 in the MS survey scan with 40.0s dynamic exclusion. The electrospray voltage applied was 2.1 kV. MS1 spectra were obtained with an AGC target of 4E5 ions and a maximum injection time of 50ms, and MS2 spectra were acquired with an AGC target of 5E4 ions and a maximum injection set to dynamic. For MS scans, the m/z scan range was 375 to 1800. The resulting MS/MS data was processed using Sequest HT search engine within Proteome Discoverer 2.5 SP1 against a Homo sapiens protein database obtained from Uniprot including the Theiler viral polyprotein and PKR-Apex sequences. Trypsin was specified as cleavage enzyme allowing up to 2 missed cleavages, 4 modifications per peptide and up to 5 charges. Mass error was set to 10 ppm for precursor ions and 0.1 Da for-fragment ions (LUMOS) or 10ppm (Exploris240). Oxidation on Met (+15.995 Da), conversion of Gln (−17.027 Da) or Glu (−18.011 Da) to pyro-Glu at the peptide N-term were considered as variable modifications. False discovery rate (FDR) was assessed using Percolator and thresholds for protein, peptide and modification site were specified at 1%. For abundance comparison, abundance ratios were calculated by Label Free Quantification (LFQ) of the precursor intensities within Proteome Discoverer 2.5 SP1.

PSM Ratio calculations were as followed:

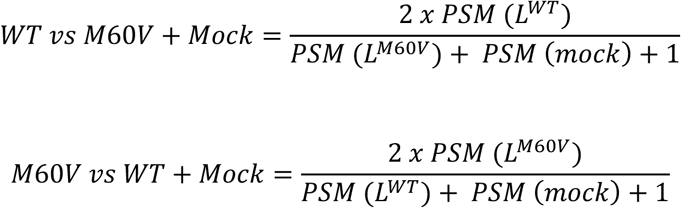

### Microscopy and Image analysis

Pictures were taken with either a spinning disk confocal microscope (Zeiss) or with an LSM980-multiphoton confocal microscope (for high-resolution microscopy). Pictures were taken with the same exposure time, image brightness and contrast. Analysis of the image was made using the Zen system image analysis software (Zeiss), the HALO image analysis platform or the CellProfiler software. Statistical analysis on immunofluorescence experiments was done using GraphPad Prism v9. One-way ANOVA with Dunnett’s multiple comparisons tests were used to compare all samples with TMEV-L^WT^. The number of independent experiments (n) and statistical comparison groups are indicated in the Figures and Figure legends. (* p< 0.05, ** p<0.01, *** p<0.001, **** p<0.0001).

For live cell imaging, HeLa PKR-S11 were plated on a 35mm glass-bottom dish (Mattek) at 1:5 confluence. Twenty-four hours later, the culture medium was replaced with a culture medium without phenol red and probe was added for 3 hours to label the DNA (SPY555-DNA, Spirochrome, 1/1000). Pictures were recorded every 10 minutes with a Zeiss apotome epifluorescence microscope equipped with an incubation chamber controlling the temperature and the CO2 levels with a 25x oil objective.

## Acknowledgments

We are grateful to Patrick Van Der Smissen for his long-standing help in confocal microscopy.

## Supporting information captions

***Fig S1.*** *Confocal microscopy images showing the distribution of RFP-NLS and GFP-NES proteins in live HeLa LVX cells infected for 10h at an MOI of 5 PFU/cell*.

***Fig S2.*** *A) Confocal microscopy images showing the co-immunostaining of SFPQ and 3D in HeLa cells infected with TMEV-L^WT^ or -L^M60V^ virus for 12h with 5 PFU/cell. Viral polymerase 3D was detected as a control of infection. Scale bar: 10µm. B) Confocal microscopy images showing the co-immunostaining of NONO and 3D in HeLa cells infected with TMEV-L^WT^ or -L^M60V^ virus for 12h at an MOI of 5 PFU/cell*.

***Fig S3.*** *A) Western blot showing the detection of PKR and its activated form (pPKR) in lysates of HeLa M, HeLa PKR-KO and HeLa PKR-APEX cells infected for 10h at an MOI of 5 PFU/cell. 3D polymerase was detected as control of infection and ß-actin as loading control. B) Western blot of proteins biotinylated by PKR-APEX in lysates and pulled down samples of TMEV infected cells. HeLa PKR-APEX cells were infected for 4, 6, 8 or 10h at an MOI of 5 PFU/cell of TMEV-L^WT^ or -L^M60V^. Biotin-phenol was added in the medium 1h before the end of the infection. After hydrogen peroxide activation, biotinylated proteins were pulled-down using streptavidin-magnetic beads. C) Confocal microscopy images showing the co-immunostaining of PRCC and VP1 in HeLa cells infected with TMEV-L^WT^ or -L^M60V^ virus for 12h with 5 PFU/cell. VP1 was detected as a control of infection. Scale bar: 10µm*.

***Table S1*** *– Viruses and expression vectors used in this study*.

